## Supplementary Figures for "Overcoming Resolution Attenuation During Tilted Cryo-EM Data Collection"

### Slide 1
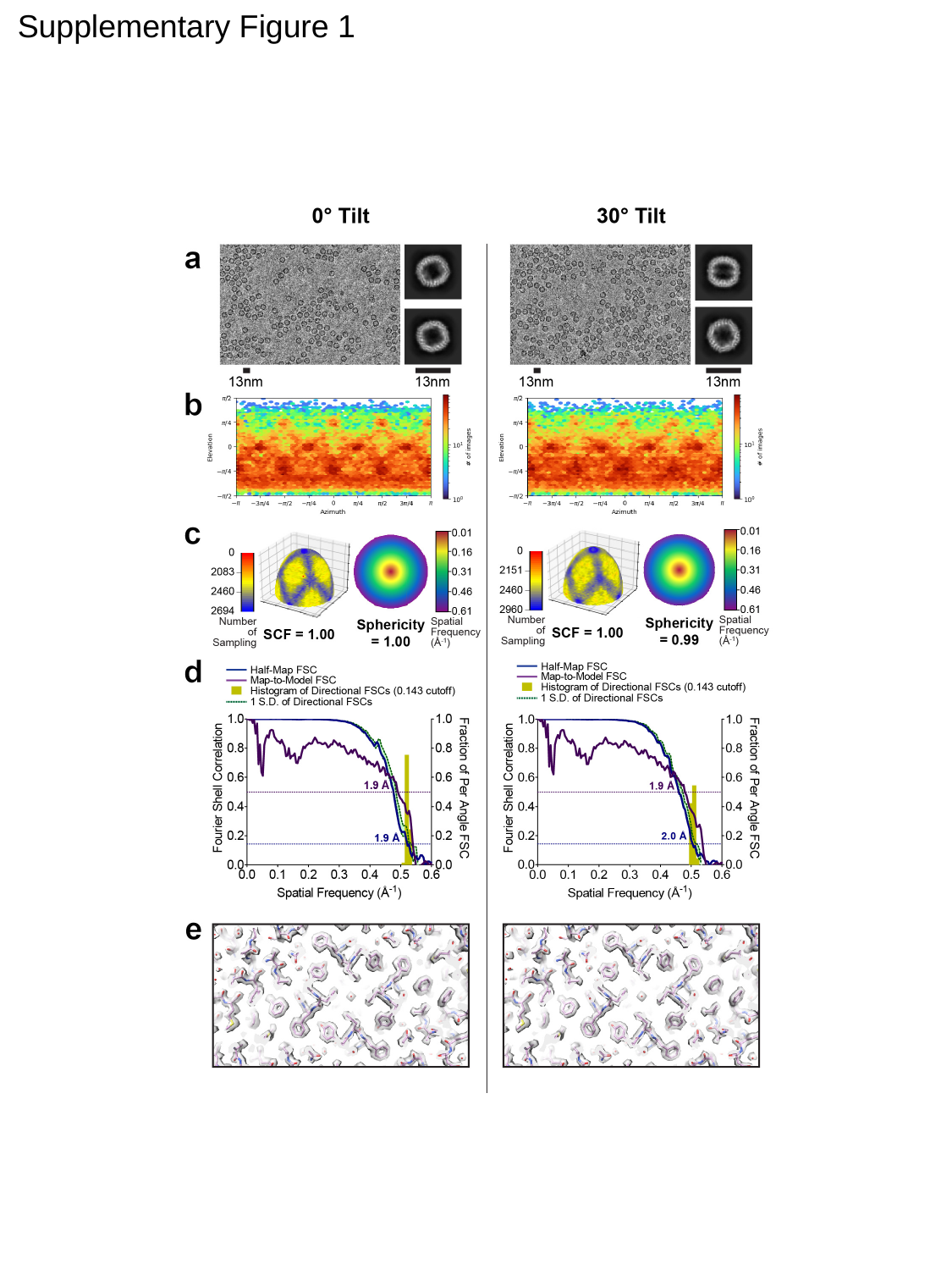

Supplementary Figure 1

### Slide 2
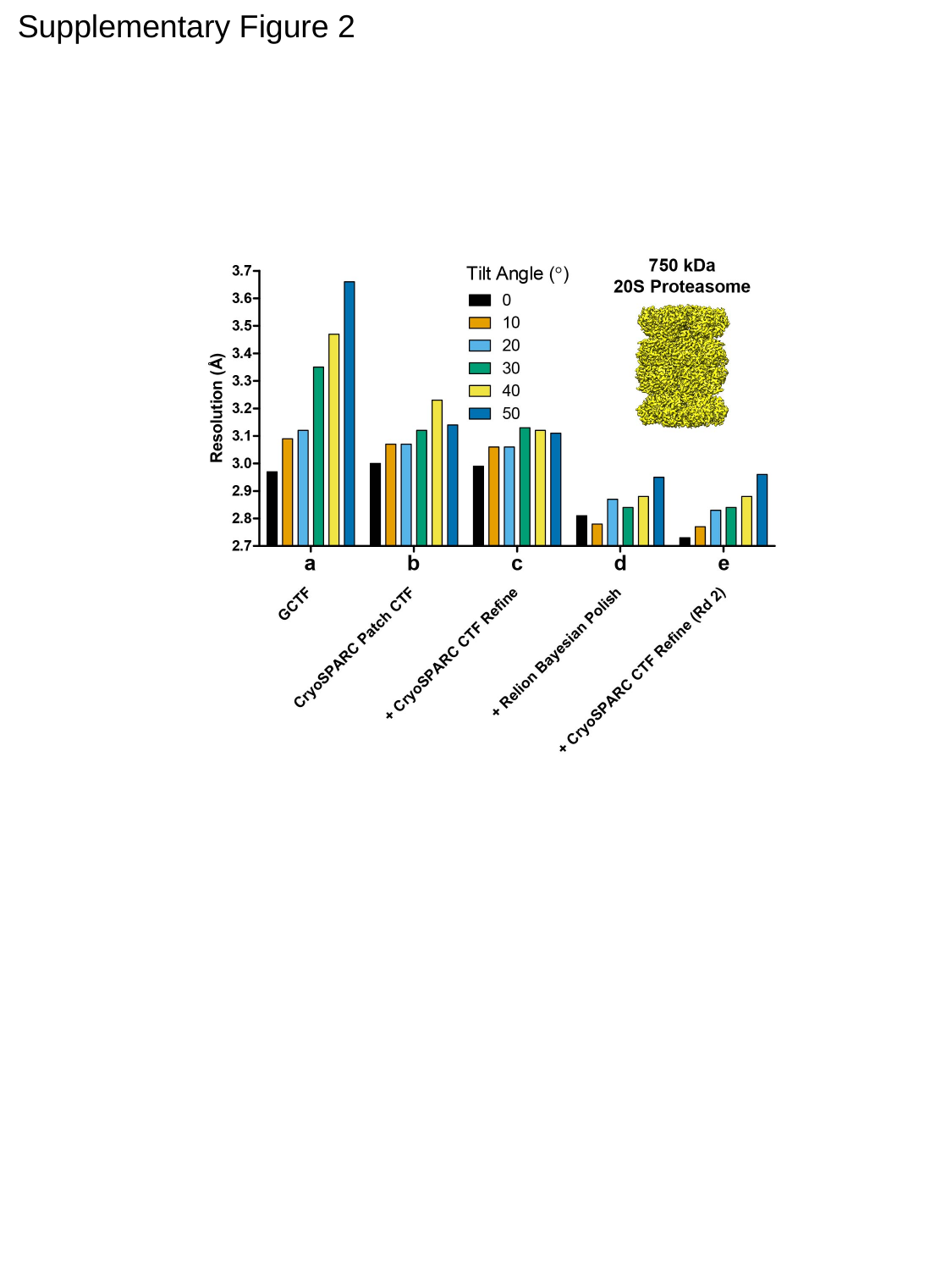

Supplementary Figure 2

### Slide 3
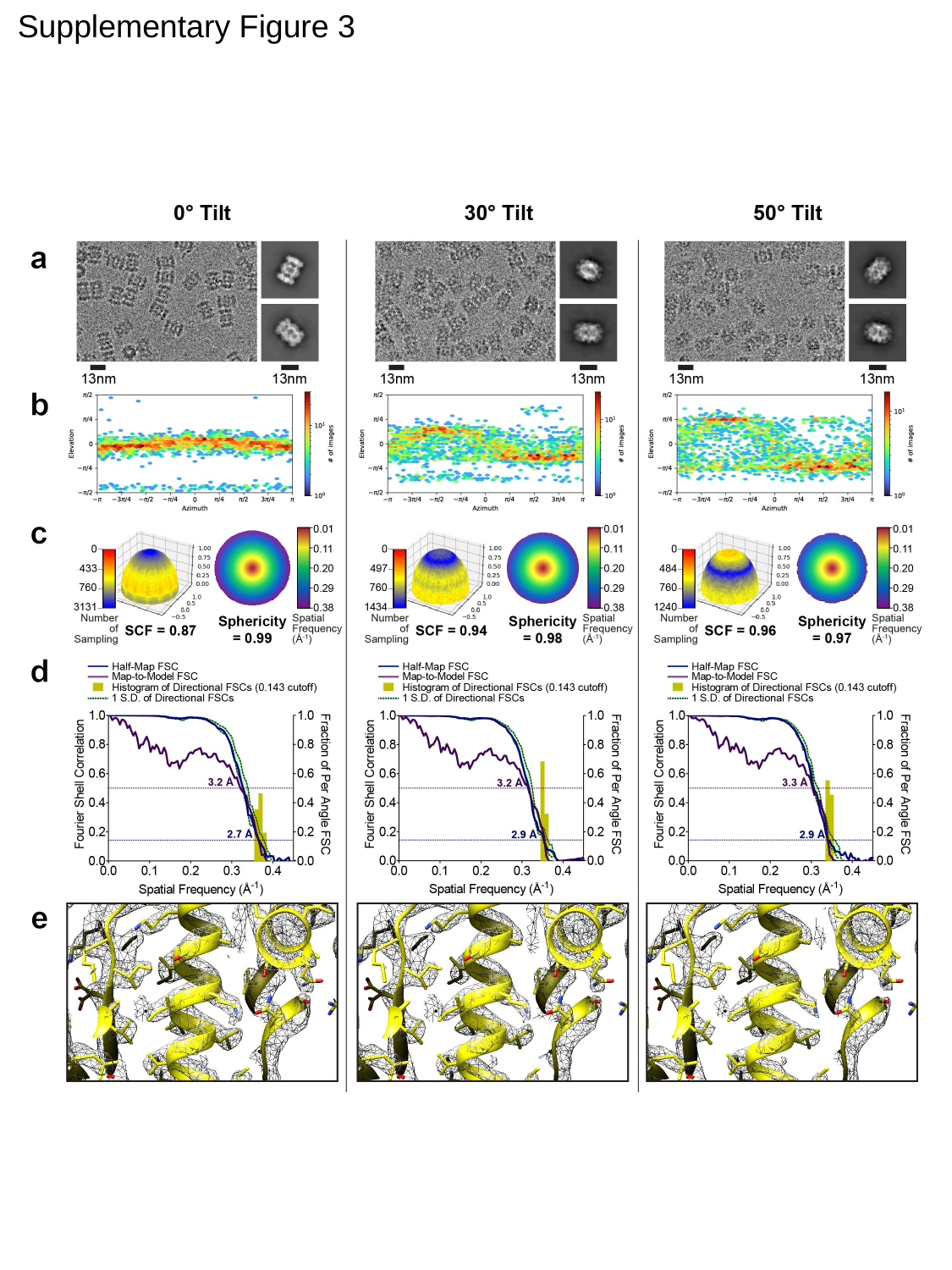

Supplementary Figure 3

### Slide 4
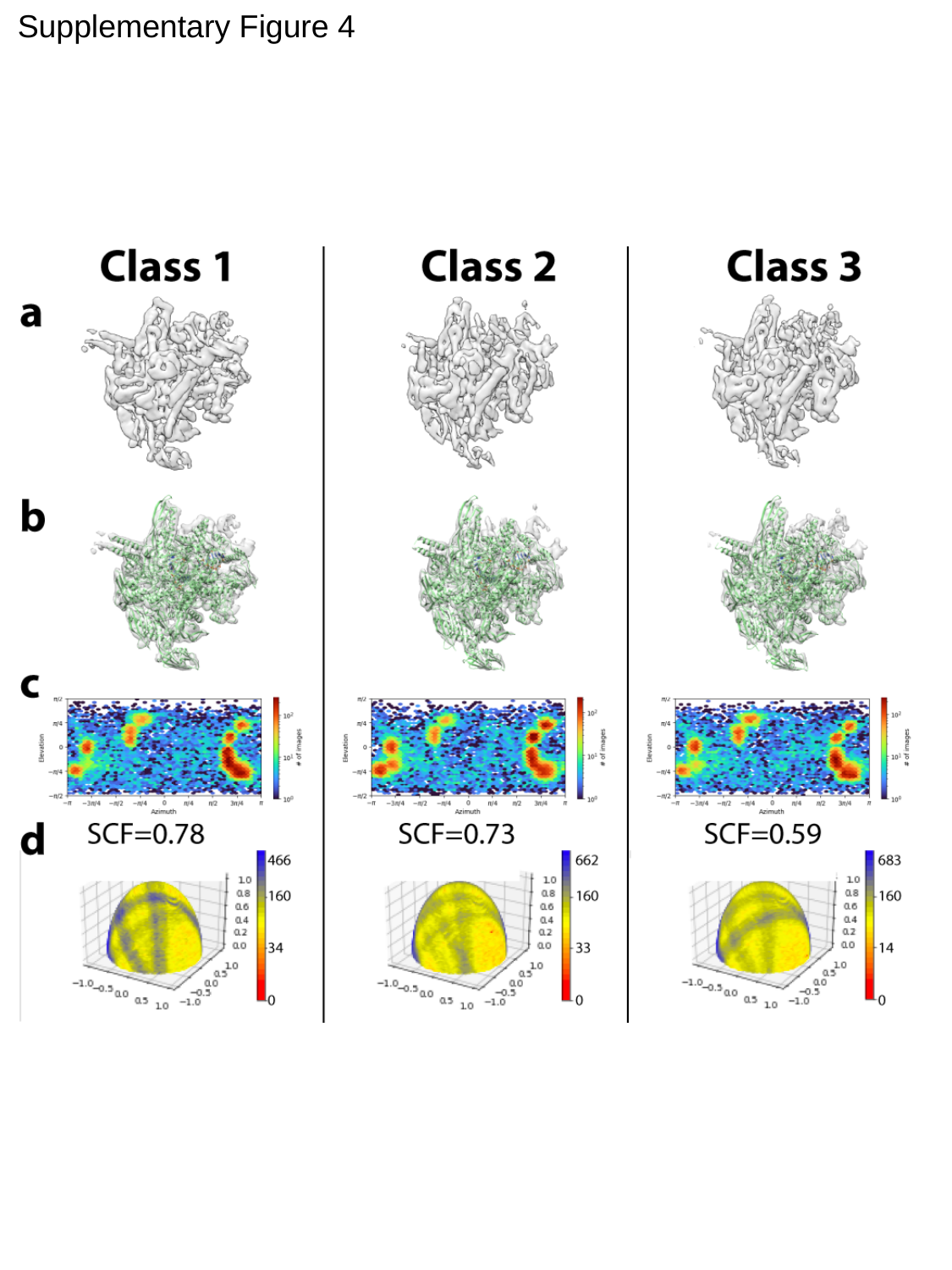

Supplementary Figure 4

### Slide 5
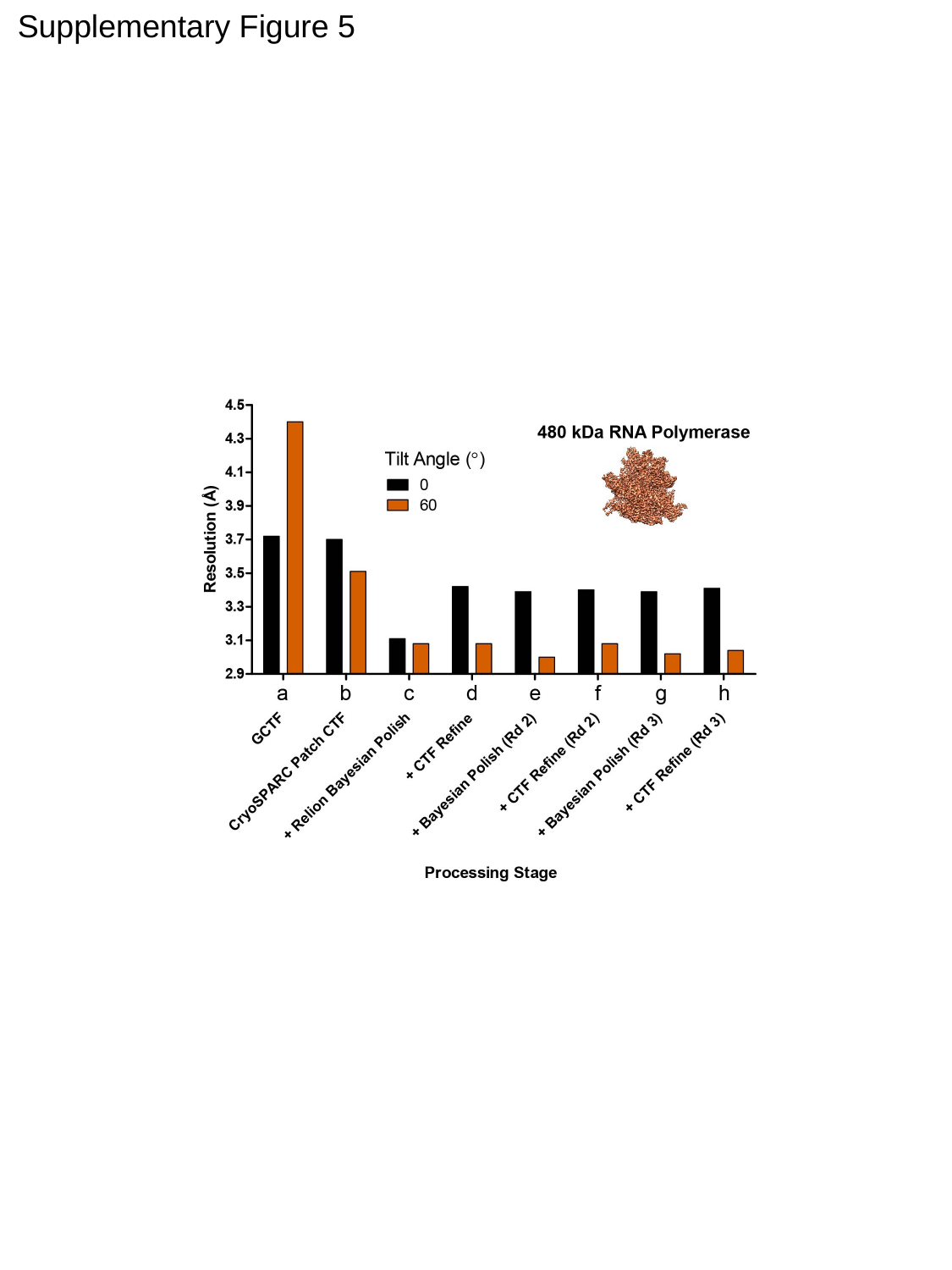

Supplementary Figure 5
